# Biogenic action of *Lactobacillus plantarum* SBT2227 promotes sleep in *Drosophila melanogaster*

**DOI:** 10.1101/2021.11.15.468562

**Authors:** Taro Ko, Hiroki Murakami, Azusa Kamikouchi, Hiroshi Ishimoto

**Author notes:** These authors contributed equally (Taro Ko, Hiroki Murakami).

## Abstract

Lactic acid bacteria (LAB) influence multiple aspects of host brain function via the production of active metabolites in the gut, which is known as the pre/pro-biotic action. However, little is known about the biogenic effects of LAB on host brain function. Here, we reported that the *Lactobacillus plantarum* SBT2227 promoted sleep in *Drosophila melanogaster*. Administration of SBT2227 primarily increased the amount of sleep and decreased sleep latency at the beginning of night-time. The sleep-promoting effects of SBT2227 were independent of the existing gut flora. Furthermore, heat treatment or mechanical crushing of SBT2227 did not suppress the sleep-promoting effects, indicative of biogenic action. Transcriptome analysis, and RNAi mini-screening for gut-derived peptide hormones revealed the requirement of neuropeptide F, a homologue of the mammalian neuropeptide Y, for the action of SBT2227. These biogenic effects of SBT2227 on the host sleep provide new insights into the interaction between the brain and gut bacteria.

## INTRODUCTION

Nutrients obtained from food sources are not only necessary for growth and sustenance but also for various vital functions and, in some cases, for improving adaptability (Milton, 2003; Watanabe et al., 2019). In general, fermentation increases the rate of nutrient absorption, increasing the nutritional value of food (Kårlund et al., 2020; Sharma et al., 2020). Animals prefer fermented food and consume the bacterial population responsible for fermentation along with it. Therefore part of the intestinal microbiota is constructed by bacteria consumed (Pasolli et al., 2020).

In mammals, intestinal bacteria are known to improve host health via their probiotic functions such as activation of immunity (Shi et al., 2017) and anti-obesity action (Barathikannan et al., 2019), and by supplying nutrients (Yadav et al., 2018), as well as via prebiotic functions that regulate the balance of intestinal bacteria (Gibson et al., 2017; Hill et al., 2014). In addition to these various biological functions, intestinal bacteria can affect higher-order brain functions, such as anxiety (Peirce and Alviña, 2019), depression (Peirce and Alviña, 2019), and cognitive functions (Chu et al., 2019), via communication with the brain–gut axis (Grenham et al., 2011; Mayer et al., 2014; Peirce and Alviña, 2019). In the past decade, the probiotic and prebiotic functions of these intestinal bacteria have attracted considerable attention, and the causative factors and mechanisms have been elucidated (Erny et al., 2015; Lyte, 2014; Strandwitz, 2018; van de Wouw et al., 2018). Furthermore, functional substrates produced by bacteria may direct biological effects irrespective of the presence of intestinal flora, a concept known as biogenics (Mitsuoka, 2000). For example, biogenic actions of bacterial substrates have been found to potentially prevent cancer (Badr El-Din et al., 2020; Zhang et al., 2013), reduce body fat (Nakamura et al., 2016), modulate lipid metabolism (Monagas et al., 2010), prevent hypertension (Cheng and Pan, 2017) and inflammation (Yang et al., 2017, 2015), and activate the immune system (Chanson-Rolle et al., 2015; Kobatake and Kabuki, 2019). Compared to probiotics and prebiotics, knowledge regarding the effects of biogenics on higher-brain functions is limited.

Recently, the fruit fly *Drosophila melanogaster*, a model organism with a simple brain and nervous system, is being increasingly used for studying the brain–gut–microbiome axis (Broderick and Lemaitre, 2012; Wong et al., 2011). The intestinal flora of the fly is a miniature version of that of mammals, with *Lactobacillus* and *Acetobacter* being the main bacterial species (Erkosar et al., 2013). Furthermore, similarities between flies and mammals have already been shown in host–microbe interactions, such as in the immune system and metabolic pathways (Buchon et al., 2014; Wong et al., 2016). Indeed, studies on the gut microbiota of *D. melanogaster* have revealed probiotic effects on brain functions such as locomotion (Schretter et al., 2018), foraging (Leitão-Gonçalves et al., 2017; Wong et al., 2017), odor-guided egg laying (Qiao et al., 2019), and food preference (Venu et al., 2014). These evidences indicated that *D. melanogaster*, with its compact brain–gut axis structure, can be used as an experimental system for studying the biogenic functions of digested bacteria using the many sophisticated molecular genetic tools that are currently available.

Sleep is one of the behaviors regulated by higher-order brain functions. Behavioral genetics of *D. melanogaster* have revealed genes and neural mechanisms that regulate sleep, most of which are shared with mammals (Hendricks et al., 2000; Ly et al., 2018; Sehgal and Mignot, 2011; Shaw, 2000). A recent study showed that the absence of commensal bacteria or axenic conditions affects sleep in flies negligibly (Jia et al., 2021; Selkrig et al., 2018). This indicates that probiotics and prebiotics in the gut microbiota do not significantly affect fly sleep, although evidence for biogenics is inconclusive so far. This gap in our knowledge of the biogenic effects could be filled by a detailed analysis of the sleep-behavior of flies following oral administration of a particular LAB strain. To achieve this, we used a *L. plantarum* strain, SBT2227 and demonstrated for the first time that the biogenic effect of the LAB can facilitate the fly sleep. Furthermore, transcriptome analysis suggested that the endocrine system is a potential target for the action of the biogenic actions. The results of a mini-screening using flies with gene knockdowns suggested the involvement of a specific peptide hormone, neuropeptide F, in the biogenic action of SBT2227. These findings provide a novel perspective on the brain-gut-microbe interaction and open up new avenues for the functional use of lactic acid bacteria.

## RESULTS

### Effects of oral administration of SBT2227 on fly sleep

*L. plantarum* is a gram-positive LAB commonly found in plants and fermented food (Guidone et al., 2014; Mundt and Hammer, 1968) and is also known to be one of the major intestinal bacteria of *D. melanogaster* (Chandler et al., 2011; Cox and Gilmore, 2007). As fruit flies may ingest *L. plantarum* from fermented fruits in the field, it is highly possible that this lactobacillus species influences the biological activities of flies. Therefore, we decided to investigate the biogenic effects of SBT2227, a strain of *L. plantarum*, on the sleep of fruit flies.

The sleep pattern of flies orally administered SBT2227 (2.6 × 10^10^ cfu/mL in food) was compared to that of untreated control flies (**Figure 1A**). Flies in both groups showed typical sleep patterns reported previously: napping in the middle of the subjective daytime and sleeping mostly during subjective night-time (Andretic and Shaw, 2005; Hendricks et al., 2000; Shaw, 2000). Daytime sleep was significantly reduced on the 1^st^ day of SBT2227 treatment, although this was not sustained after the 2^nd^ day (**Figure 1B**). By contrast, obvious change was not detected in night-time sleep on the 1^st^ day, although it increased significantly on the 2^nd^ and 3^rd^ day in SBT2227-fed flies (**Figure 1C**).

**Figure 1.**
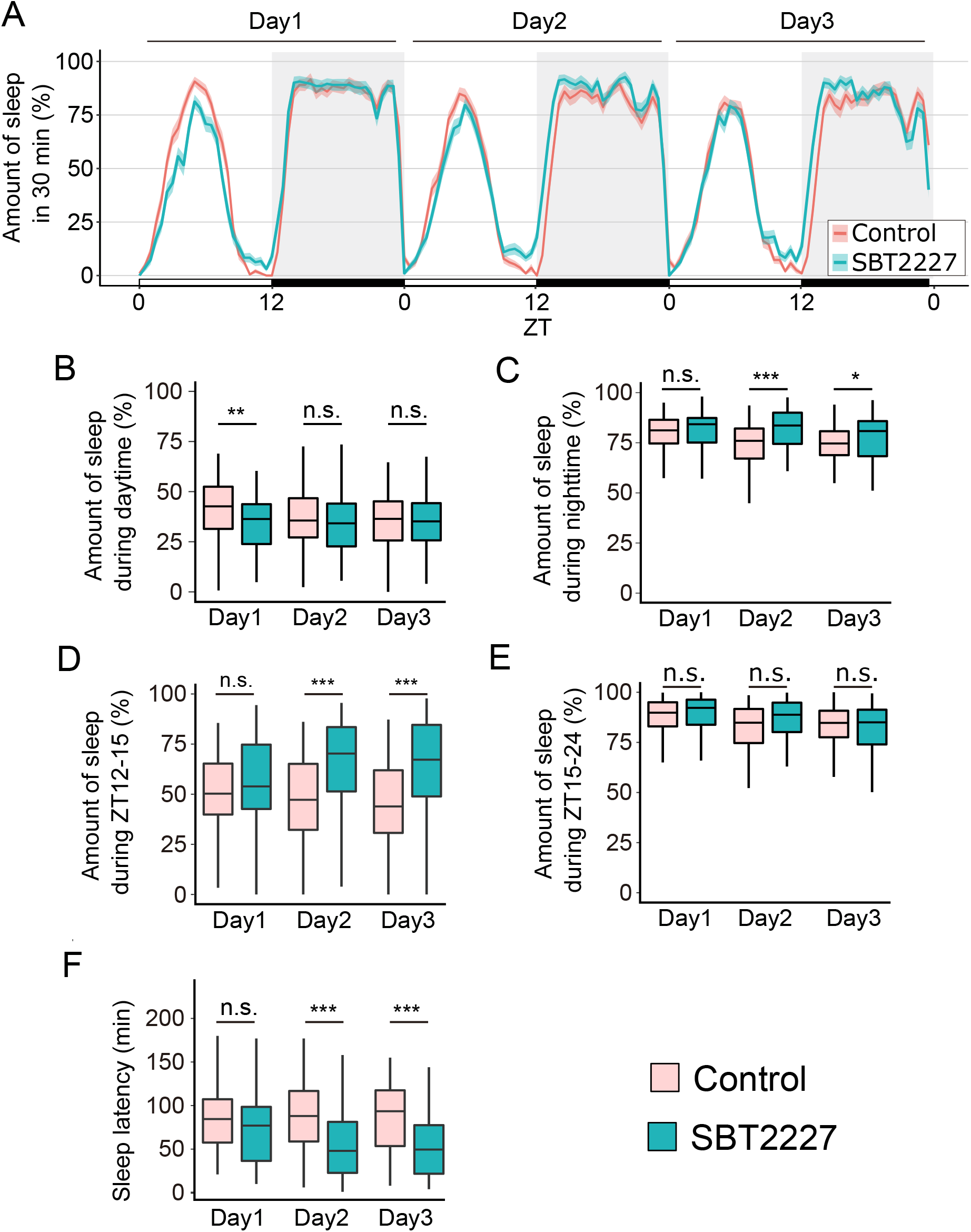
Oral administration of SBT2227 increased sleep at the onset of night-time and decreased sleep latency. (A) Sleep patterns of flies fed the control food (red) or SBT2227 food (green). Sleep traces are presented as mean ± SEM. (B, C) Amount of sleep during daytime (ZT0-12) (B) and night-time (ZT12-24) (C). (D, E) Amount of sleep at specific timing; ZT12-15 (D) and ZT15-24 (E) are shown. (F) Sleep latency. n = 96 for each group. Wilcoxon-Mann-Whitney test was used for statistical analysis. * *p* < 0.05, ** *p* < 0.01; *** *p* < 0.001, n.s.; not significant. Red, flies fed the control food; Green, flies fed SBT2227 food.

Based on the night-time sleep pattern on day 3 (**Figure 1A**), the sleep-promoting effect was preferentially detected at a specific time window (ZT12-15, Zeitgeber time (ZT), where ZT0 = lights on and ZT12 = lights off, **Figure 1D**). By contrast, no obvious sleep effect was detected during ZT15-24 (**Figure 1E**). The sleep effect appeared at early night-time, presumably because of the shorter latency of sleep. Sleep latency was significantly reduced (**Figure 1F**). These effects of SBT2227 (increase in sleep amount during ZT12-15 and shortened sleep latency on the 3^rd^ day) were also conserved in another fly strain with a different genetic background (**Figure S1**). Therefore, we decided to use these two characteristics, sleep amount of ZT12-15 and sleep latency on the 3^rd^ day, as the main parameters for further analyses, as they are considered to show the typical effects of SBT2227.

### Effects of oral administration of SBT2227 on awakening state

Sleep and wakefulness are two sides of the same coin; as sleep increases, wakefulness decreases, and vice versa. However, this reciprocal relationship is not always true. For example, Fernandez-Chiappe *et al*. reported a case where flies did not increase their sleep when they reduced the activity level (Fernandez-Chiappe et al., 2020). On the other hand, Potdar and Sheeba reported another case where flies increased sleep without changing their activity level (Potdar and Sheeba, 2018). Therefore, we investigated the effects of SBT2227 on wakefulness. The activity of SBT2227*-*fed flies was examined over a 3-day period. The graph showing the activity patterns showed that the flies fed SBT2227 showed a clear decrease in their night-time activity after the second day (**Figure 2A**). Separate quantitative analysis of daytime and night-time activity counts showed absence of any obvious change in the daytime; however, a significant decrease was detected at night-time in flies fed SBT2227 (**Figure 2B, C**). Reduction in activity counts may be attributed to the decrease in motor control. To assess whether SBT2227 disrupts motor ability, we examined the activity index, which is the number of activity counts per waking time. We did not detect any obvious change in the activity index during the daytime, which is the main waking period of flies (**Figure 2D)**. By contrast, we found a significant reduction in the activity index at night on day 3 (**Figure 2E)**. As daytime locomotion was normal in flies fed SBT2227, the night-time arousal level on the 3^rd^ day was lower than that of the control; in other words, sleep was strongly promoted.

**Figure 2.**
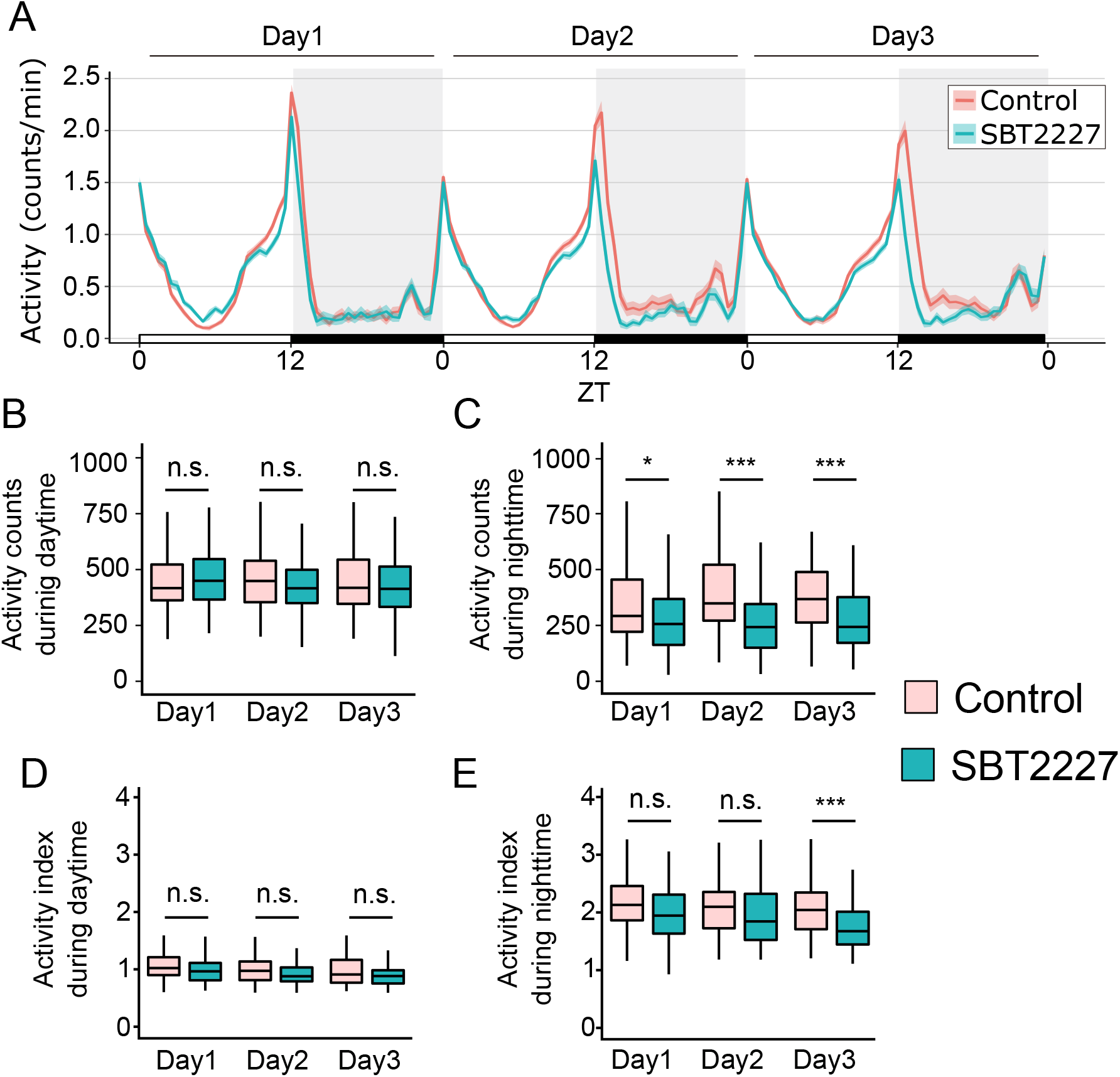
Wakefulness was altered in flies fed SBT2227. (A) Activity patterns of flies fed the control food (red) or SBT2227 food (green). (B, C) Activity counts during daytime (ZT0-12) (B), and night-time (ZT12-24) (C). (D, E) Activity index during daytime (D), and night-time (E). n = 96 for each group. Wilcoxon-Mann-Whitney test was used for statistical analysis. * *p* < 0.05, ** *p* < 0.01; *** *p* < 0.001, n.s.; not significant.

### Commensal bacteria did not alter the sleep-promoting effect of SBT2227

To investigate whether the sleep-promoting effect of SBT2227 requires interaction with the commensal bacterial flora, the existing intestinal bacteria were disturbed with antibiotics (Storelli et al., 2018) and sleep behavior was evaluated with or without SBT2227 treatment (**Figure 3A**). The number of aerobic bacteria in the <u>a</u>nti<u>b</u>iotic-treated flies (ABT flies) was reduced to approximately 1/100 compared to that in flies fed the <u>c</u>ontrol <u>c</u>onventional food (CC flies) (**Figure 3B**). LAB, including *L. plantarum*, are anaerobic bacteria. Therefore, we determined the number of anaerobic bacteria present in the gut and found that anaerobic bacteria were almost eliminated in ABT flies (**Figure 3B**). Significant differences were not observed between the sleeping behaviors of ABT and CC flies (**Figure 3C, D**). By contrast, the sleep-promoting effect of SBT2227 was observed in both ABT and CC flies (**Figure 3E, F**). The effect sizes of SBT2227 treatment on the sleep amount (ZT12-15) and sleep latency in CC and ABT flies were similar (**Figure 3E, F**, sleep amount effect size (*r)* = 0.35 and 0.29, and sleep latency effect size (*r)* = 0.31 and 0.34, respectively). These results suggested that intestinal commensal bacterial flora is not required for the action of SBT2227 on fly sleep.

**Figure 3.**
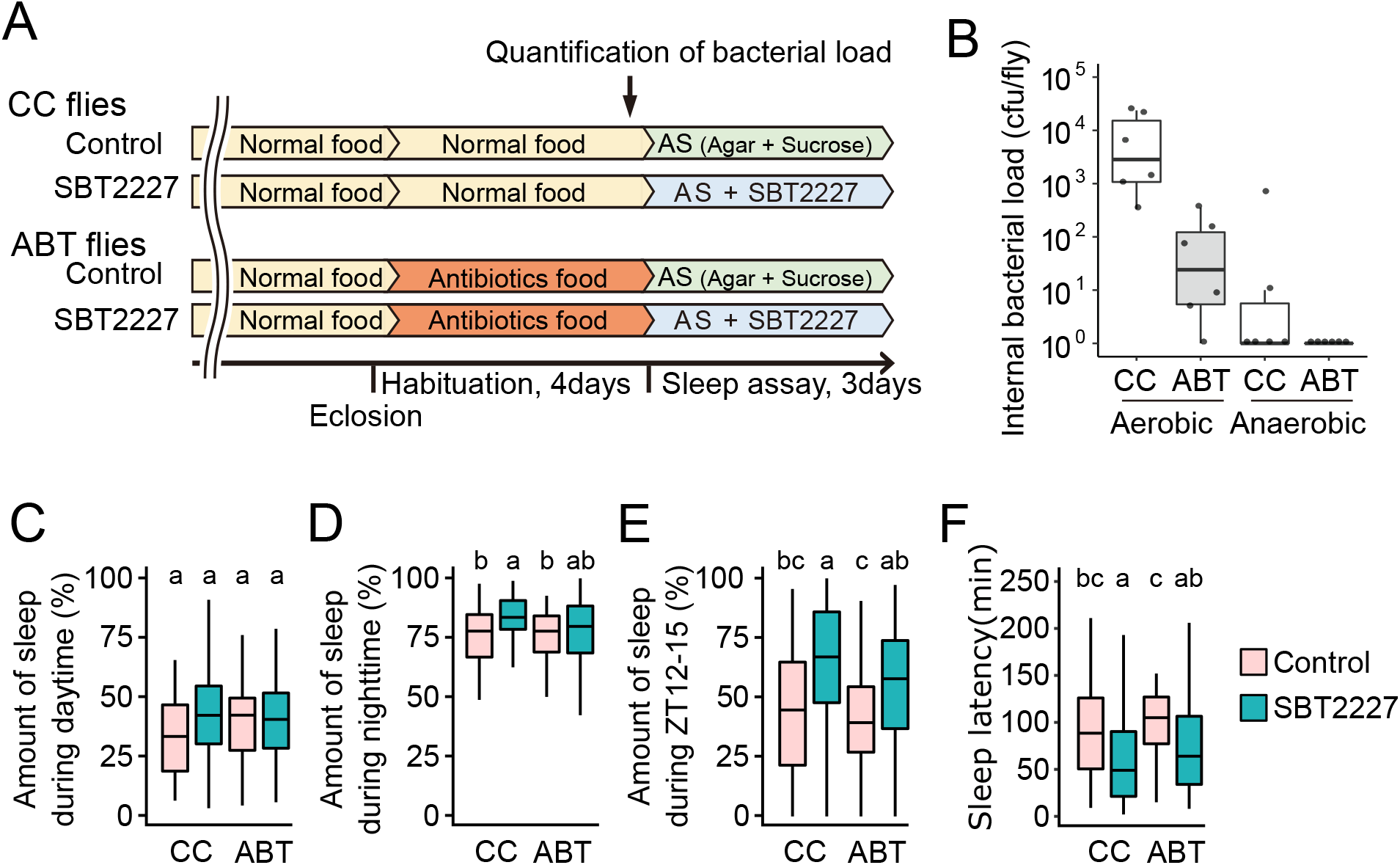
Elimination of gut microbes did not alter the sleep effects of SBT2227. Flies fed the control conventional food (CC flies) or antibiotic-containing food (ABT flies) were tested for sleep. (A) Schematic diagram illustrating the experimental flow. (B) Internal bacterial load in CC and ABT flies (n = 6). (C-F) Amount of sleep during daytime on day 3 (C), amount of sleep during night-time on day 3 (D), amount of sleep during ZT12-15 on day 3 (E), and sleep latency on day 3 (F). (C−E) n ≥ 58 for each group. The Steel-Dwass-Critchlow-Fligner method was used for statistical analysis. Different letters indicate statistical differences between groups (*p* < 0.05).

### The sleep-promoting effect of SBT2227 was sustained after protein denaturation

In order to clarify whether the SBT2227 needs to be alive to have the sleep-promoting effect, heat-killed SBT2227 was administrated to flies and evaluated its effect on sleep. SBT2227 was treated with 65°C for 1 h or autoclaved at 121°C for 15 min and then administered to flies. Compared to that in the control, the amount of sleep during ZT12-15 on day 3 increased significantly in all the SBT2227-fed groups (unheated, 65°C, or autoclaved) (**Figure 4A**). Similarly, compared to that of the control group, sleep latency decreased significantly in all SBT2227 treated groups (**Figure 4B**). These results suggested that SBT2227 does not need to be alive for the sleep-promoting effect. Moreover, protein denaturation in SBT2227 did not alter its sleep-promoting effect. The heat-stable substances in SBT2227 were possibly the biogenic factors required for the sleep-promoting effect.

**Figure 4.**
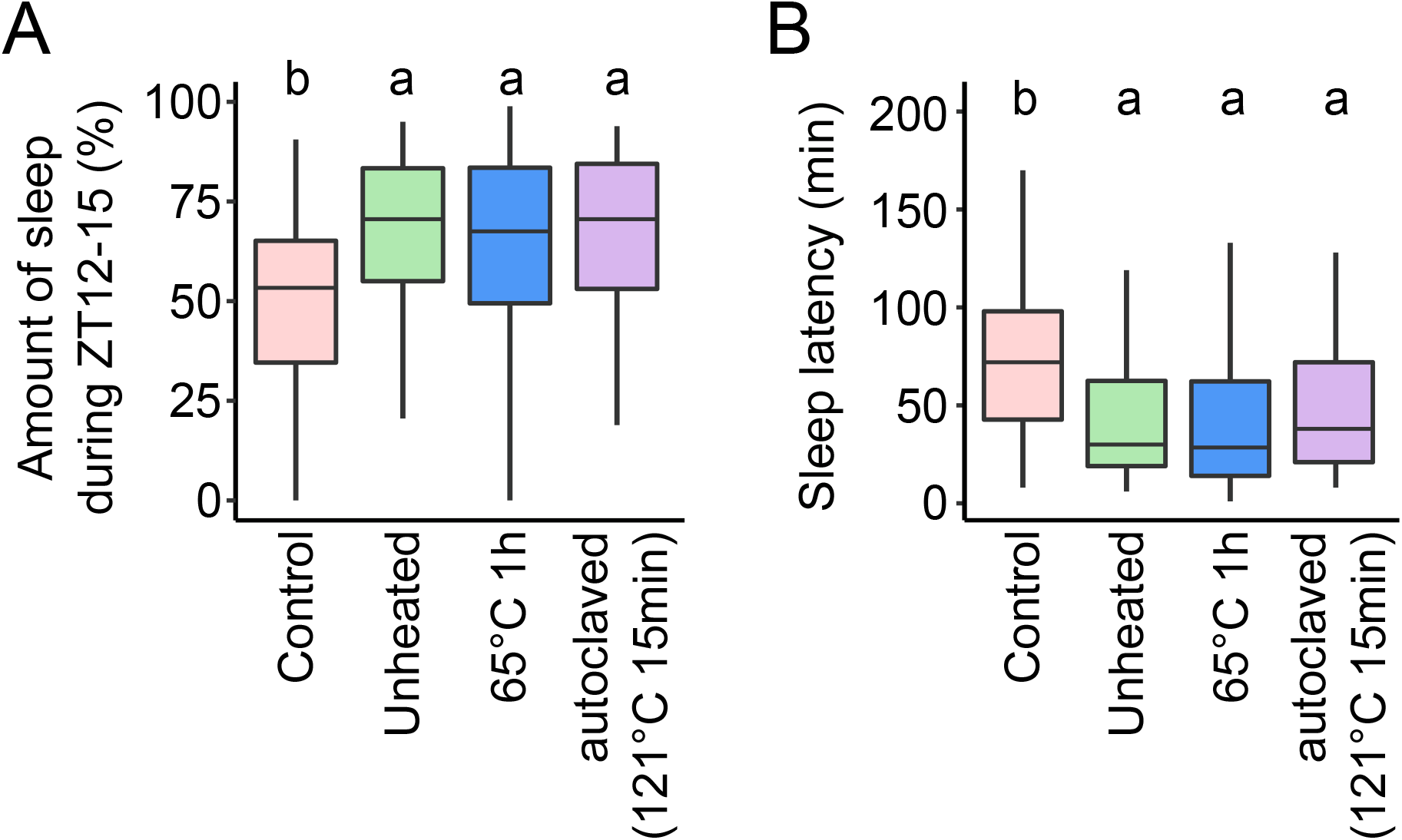
Heat denaturation of SBT2227 did not alter its effects on fly sleep. (A, B) Amount of sleep during ZT12-15 on day 3 (A), and sleep latency on day 3 (B) of flies fed the control food (red), unheated SBT2227 (green), heat-treated SBT2227 (65°C for 1 h, blue), or autoclaved SBT2227 (121°C for 15 min, purple). n = 95−96 for each group. The Steel-Dwass-Critchlow-Fligner method was used for statistical analysis. Different letters indicate statistical differences between groups (*p* < 0.05).

### Supernatant of crushed SBT2227 cells may contain active substances required for the sleep-promoting effect

To identify the SBT2227-derived components that exert biogenic effects on sleep, we crushed the bacteria and separated it into a supernatant rich in intracellular/membrane components and a precipitate rich in cell wall components. To verify the efficiency of crushing the bacteria, we performed Gram staining of the bacterial samples before and after crushing. Most of the bacteria were decolorized after crushing, and the cell walls were destroyed **(Figure 5A)**. We then examined the sleep effects of SBT2227 under four different conditions (uncrushed, crushed, crushed supernatant, and crushed precipitate). The results showed that irrespective of the conditions of SBT2227, the amount of sleep in ZT12-15 on day 3 increased significantly, except in the case of the crushed precipitate (**Figure 5B**). Consistently, sleep latency decreased significantly in flies treated with uncrushed, crushed, or crushed supernatants of SBT2227, except for those treated with the precipitated fraction (**Figure 5C**). These results suggested that the intracellular/membrane components of SBT2227 may contain active substances required for the sleep-promoting effects.

**Figure 5.**
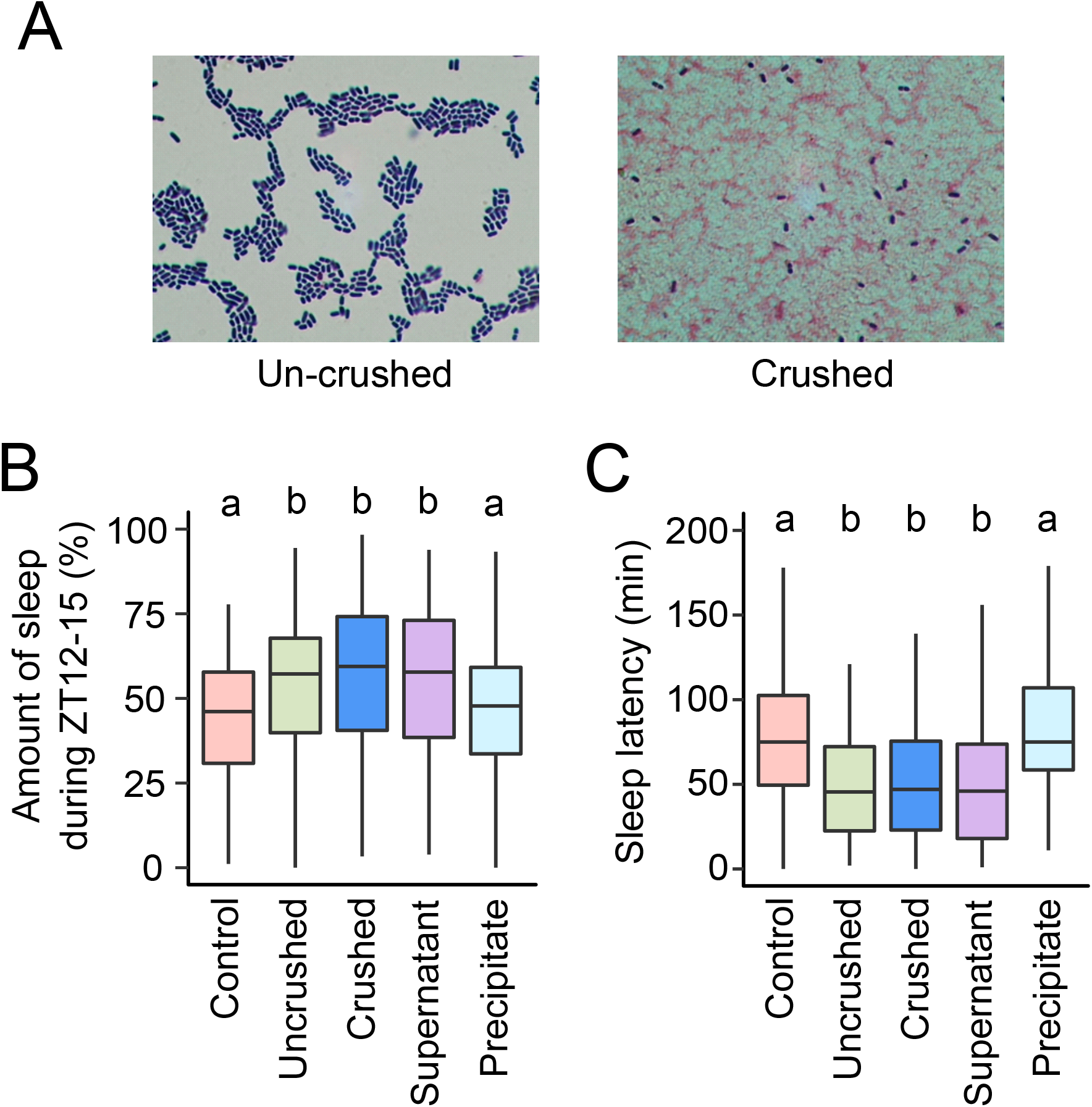
Supernatant of crushed SBT2227 promoted sleep similar to that induced by intact SBT2227. (A) Gram staining of the SBT2227 samples before and after crushing. Violet cells represent uncrushed cells. (B, C) Amount of sleep during ZT12-15 on day 3 (B) and sleep latency on day 3 (C). n = 94−96 for each group. The Steel-Dwass-Critchlow-Fligner method was used for statistical analysis. Different letters indicate statistical differences between groups (*p* < 0.05).

### Biological processes altered by SBT2227 administration

The gut is possibly a primary target of the action of the sleep-promoting substances of SBT2227, as it has abundant immune responsive cells and enteroendocrine cells (EECs), and the nerve fibers connecting the brain or ventral nerve cord to the gut have been identified (Miguel-Aliaga et al., 2018). Hence, it is possible that the substances in SBT2227 are digested and absorbed and directly affect the brain, but it is also possible that the active substances or digested and absorbed substances act on the gut, and changes that occur in the gut affect the brain via immune, endocrine, and neural pathways. Therefore, we investigated the changes in gene expression in the gut with or without SBT2227 administration using RNA sequencing-based transcriptome analysis.

The intestines containing fore-, mid-, and hindgut of flies were harvested at ZT12-14, which acts as the main time window for the sleep-promoting effect, and the mRNA extracted was subjected to RNA sequencing. The number of genes with more than one read detected in either the SBT2227-fed or SBT2227-unfed (control) group was 12,015 genes, of which 787 genes were identified as differentially expressed genes (DEGs) (**Figure 6A, see also Table S1**). Next, we conducted Gene Ontology (GO) enrichment analysis to identify the categories of molecular functions associated with these 787 genes. Multiple categories of significantly enriched DEGs were detected, including those associated with nucleosome/DNA binding, sodium ion transporter activity, neuropeptide receptor activity, and peptidoglycan muralytic activity (**Figure 6B**). Studies have shown that peptide hormones secreted by endocrine cells in the gut, such as CCHa1, act on brain neurons and regulate fly behaviors such as wakefulness (Titos and Rogulja, 2020). Therefore, among these GO categories, we focused on neuropeptide receptor activity, and hypothesized that peptide hormones produced by EECs may mediate the sleep-promoting effects of SBT2227 and tested this possibility in subsequent experiments.

**Figure 6.**
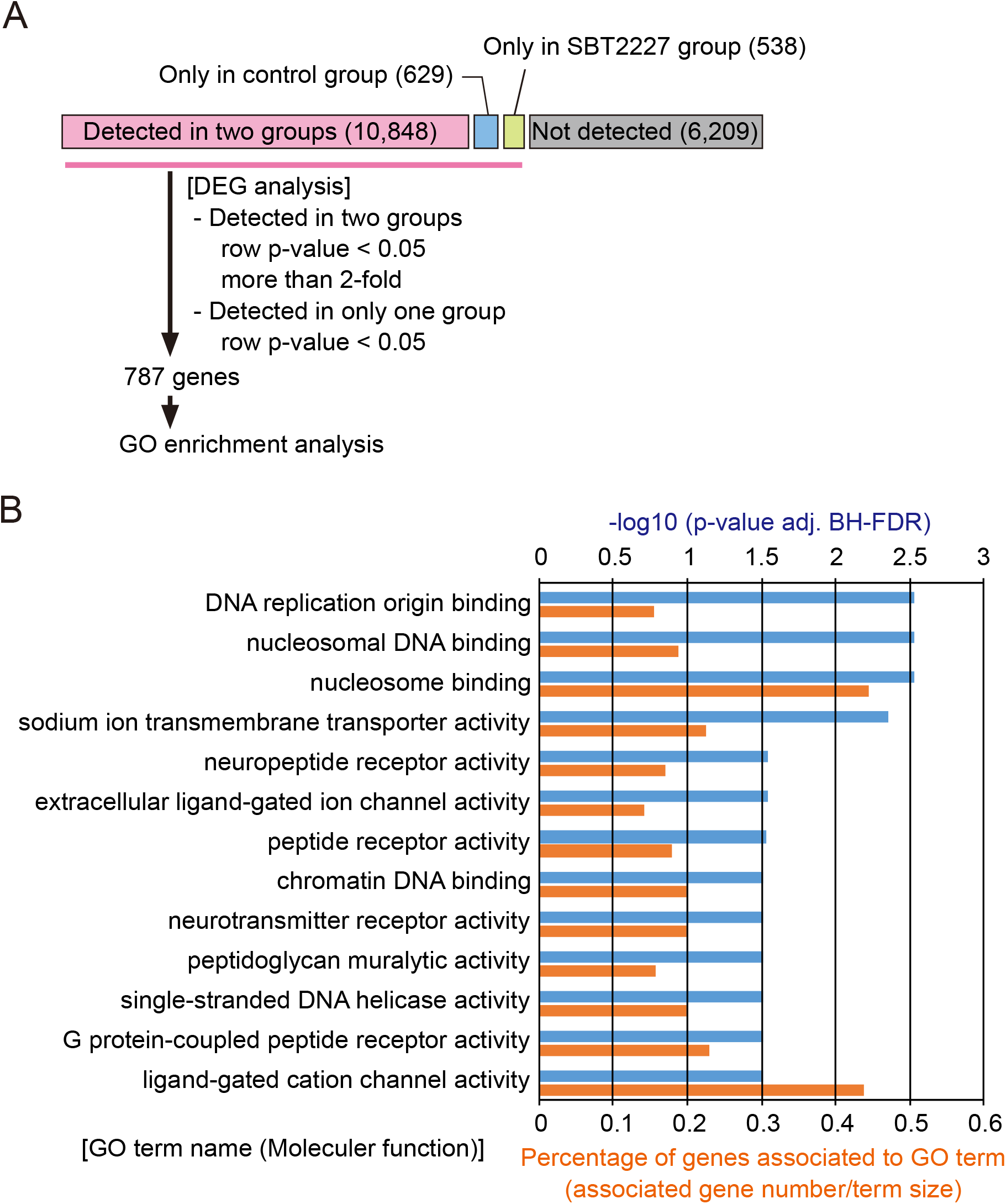
RNA-seq based transcriptome analysis indicated multiple biological pathways altered by the administration of SBT2227. (A) A flow chart of transcriptome analysis. (B) Statistically significant GO terms in category of “Molecular Function” are shown with *p*-value and enrichment score. Enrichment score was calculated as (number of genes associated with the GO term) / (number of all genes in the GO term).

### Neuropeptide F was required for the sleep-promoting effect of SBT2227

In total, 16 different peptide hormones have been reported to be expressed in EECs of *D. melanogaster* (Ji Chen et al., 2016). The expression of seven of these peptide hormones in the gut EECs, namely, allatostatin A (AstA), myoinhibiting peptide precursor (Mip), allatostatin C (AstC), CCHamide-1 (CCHa1), CCHamide-2 (CCHa2), diuretic hormone 31 (Dh31), and neuropeptide F (NPF), has been confirmed using *in situ* hybridization and immunostaining (Ji Chen et al., 2016). These hormones have also been found to be related to sleep, wakefulness, and circadian rhythms (Jiangtian Chen et al., 2016; Chung et al., 2017; Díaz et al., 2019; Fujiwara et al., 2018; He et al., 2013; Hermann et al., 2012; Kunst et al., 2014; Oh et al., 2014; Ren et al., 2015). Therefore, we focused on these seven peptide hormones and evaluated their involvement in the sleep effects of SBT2227.

First, we combined *Act-Gal4*^*25FO1*^ with *UAS-RNAi* against each target gene to ubiquitously knockdown their expression and analyzed the sleep of flies with or without administration of SBT2227. As *Act-Gal4*^*25FO1*^ *>AstA*^*RNAi*^ flies showed severe lethality, we could not evaluate them under these conditions. Quantitative comparison of the amount of sleep during the ZT12-15 period on the 3^rd^ day showed significant increase in sleep in the following order: *Act-Gal4*^*25FO1*^ *>AstC*^*RNAi*^, *Act-Gal4>CCHa2*^*RNAi*^, or *Act-Gal4*^*25FO1*^ *>Dh31*^*RNAi*^ flies fed SBT2227 (**Figure 7A**). By contrast, in *Act-Gal4*^*25FO1*^ *>Mip*^*RNAi*^, *Act-Gal4*^*25FO1*^ *>CCHa1*^*RNAi*^, or *Act-Gal4*^*25FO1*^ *>NPF*^*RNAi*^ flies, the amount of sleep did not increase after SBT2227 feeding (**Figure 7A**). Furthermore, the sleep latency decreased significantly after SBT2227 feeding in *AstC* or *CCHa2* knockdown flies, but not in *Mip, CCHa1, Dh31*, or *NPF* knockdown flies **(Figure 7B**). These results indicate that *Mip, CCHa1*, and *NPF* are required for both sleep effects (increasing the amount and decreasing the latency) of SBT2227. Thus, they are the putative candidate mediators.

**Figure 7.**
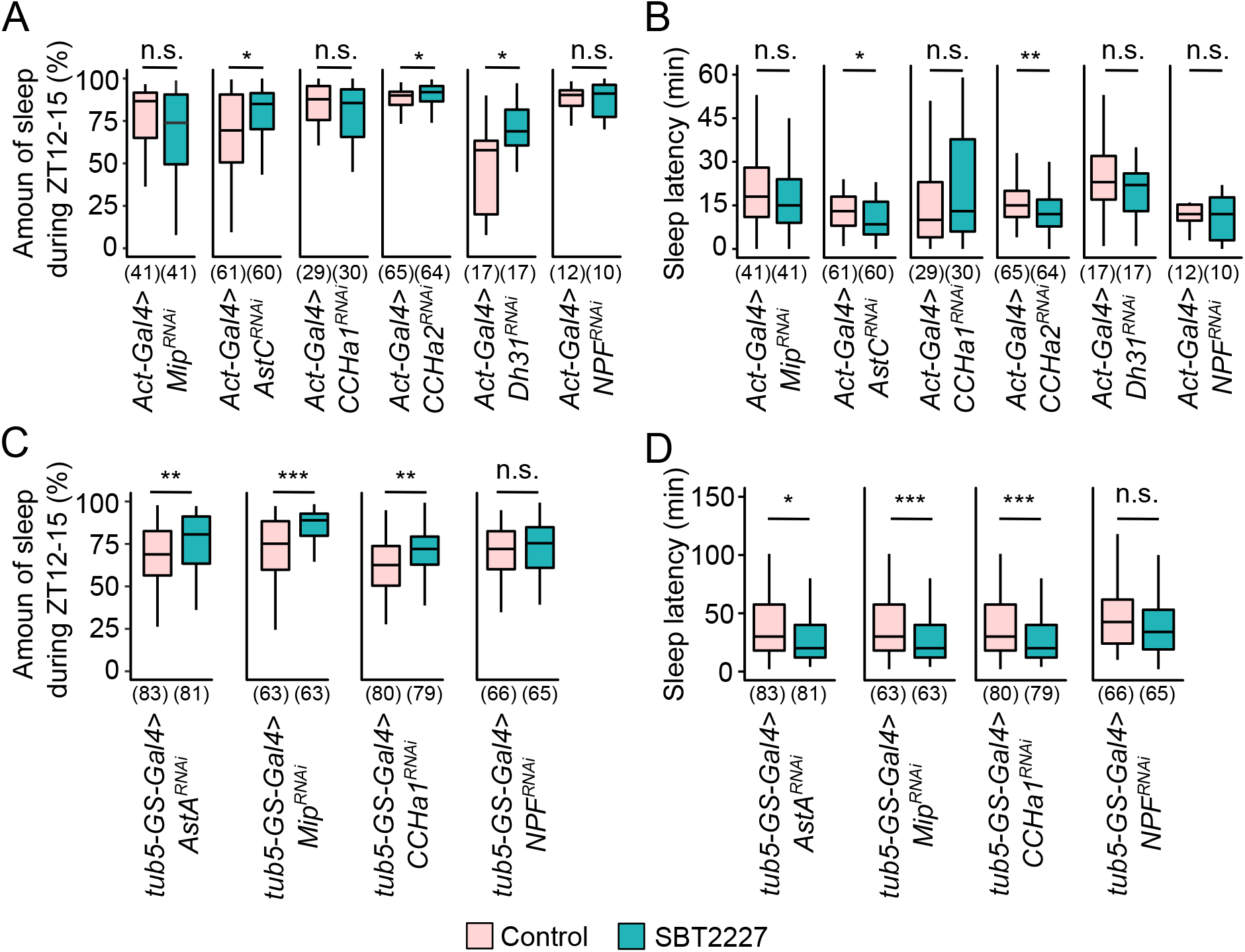
Mini-screening of flies with gene knockdown of the peptide hormones expressed in the gut. (A, B) Knockdown was performed ubiquitously using *Act-Gal4*. (A) Amount of sleep during ZT12-15 on day 3. (B) Sleep latency on day 3. (C, D) Knockdown was performed ubiquitously and temporally using *tub5-GS-Gal4*. Amount of sleep during ZT12-15 on day 3 (C) and sleep latency on day 3 (D). The sample size is shown at the bottom of each graph. The Wilcoxon-Mann-Whitney test was used for statistical analysis of control group vs SBT2227 administered group in each genotype. * *p* < 0.05, ** *p* < 0.01, *** *p* < 0.001, n.s. means not significant at *p* < 0.05.

Continuous repression of *AstA* expression using *Act-Gal4*^*25FO1*^ may affect developmental processes. To exclude the possibility of a developmental abnormality, we performed molecular genetic manipulations to temporally suppress the expression of *AstA, Mip, CCHa1*, or *NPF* using the GeneSwitch system (Osterwalder et al., 2001; Roman et al., 2001). To suppress the expression of these genes ubiquitously at the adult stage, we used *tub-GeneSwitch-Gal4* in combination with *UAS-RNAi* corresponding to each gene. Significant reduction in gene expression was observed in flies after RNAi induction with RU486 feeding (**Figure S2**). The amount of sleep during ZT12-15 on day 3 increased significantly in flies with temporary knockdown of *AstA, Mip*, or *CCHa1*, but not in flies with *NPF* knockdown (**Figure 7c**). In addition, the effect of SBT2227 administration on sleep latency was not observed in flies with *NPF* knockdown **(Figure 7d**). These results suggested that *NPF* was the most potent mediator of the sleep-promoting effects of SBT2227.

## DISCUSSION

We investigated the biological effects of LAB feeding using the fruit fly *D. melanogaster* and found the sleep-promoting effects of the LAB strain *L. plantarum* SBT2227. SBT2227 retained its sleep-promoting effects even after heat denaturation. Therefore, SBT2227 did not act on the host organism as a probiotic. Furthermore, SBT2227 promoted sleep in flies even when commensal intestinal bacteria were removed, suggesting that it did not act as a prebiotic. This finding is consistent with that of a previous study showing that removal of the *Drosophila* intestinal flora negligibly affects sleep (Jia et al., 2021; Storelli et al., 2018). These results suggested that the biological action of SBT2227 is mostly biogenic. In biogenic action, substances present in LAB induce biological effects (Mitsuoka, 2000). The supernatant of the cytoreductive fluid of SBT2227 promoted sleep in flies, suggesting that the active substances were present in the supernatant fraction. The fact that autoclaved SBT2227 promoted sleep in flies suggested that heat-denatured proteins and hydrolyzed DNA/RNA can be excluded from the list of potential active substances. Molecules such as peptidoglycans and cell wall-associated components in the precipitated fraction of crushed SBT2227 may also be excluded as candidates, as they did not exert any sleep-promoting effect under our study conditions. The active substances can be identified by comprehensively identifying the substances in the supernatant fraction and administering the identified substances to the flies.

RNA-seq-based gut transcriptome analysis in this study revealed multiple biological pathways that were enriched by SBT2227 administration, which may contain molecules responsible for SBT2227’s action on fly sleep. Neuroactive peptide hormones are involved in these candidate biological pathways. The first candidate for biogenic action of LAB in the target tissue is the cell population comprising the intestine, as it is the site of absorption of the active substance of LAB. EECs are one such group of cells (Ji Chen et al., 2016; Guo et al., 2019; Hung et al., 2020), from which peptide hormones are secreted into the hemolymph, which possibly transfers these hormones to brain regions to regulate fly sleep. Some EEC-derived peptide hormones, such as *AstA, Mip, AstC, CCHa1, CCHa2, Dh31*, and *NPF*, have been reported as sleep/wake-affective or circadian rhythm-affective hormones (Jiangtian Chen et al., 2016; Chung et al., 2017; Díaz et al., 2019; Fujiwara et al., 2018; He et al., 2013; Hermann et al., 2012; Kunst et al., 2014; Oh et al., 2014; Ren et al., 2015). To identify the molecules responsible for the sleep-promoting effect of SBT2227 from among these peptide hormones, we performed mini-screening for suppression of gene expression by combining RNA interference (Fire et al., 1998) with *Gal4/UAS* method (Fischer et al., 1988). NPF was found to be an endocrine factor necessary for the sleep-promoting effect of SBT2227.

NPF is a peptide hormone composed of 36 amino acid residues, the vertebrate homolog of which is neuropeptide Y (NPY) (Brown et al., 1999). NPFR, the receptor for NPF, encodes a G-protein coupled receptor (GPCR)-type receptor that activates inhibitory G proteins (Garczynski et al., 2002). The biological functions of NPF include regulation of feeding (Wu et al., 2005), courtship behavior (Liu et al., 2019), alcohol sensitivity (Wen et al., 2005), aggression (Dierick and Greenspan, 2007), and learning and memory (Krashes et al., 2009). It is also involved in the regulation of circadian rhythms, especially in the evening (Hermann et al., 2012). In zebrafish, NPY promotes sleep by inhibiting the wake-promoting noradrenergic system (Singh et al., 2017). By contrast, the sleep-promoting effect of NPY in mammals is still controversial; intravenous injection of NPY promotes sleep in humans (Antonijevic et al., 2000; Held et al., 2006). Furthermore, in rodents, injection of NPY into the brain has been reported to promote sleep and decrease locomotion (Akanmu et al., 2006; Jolicoeur et al., 1991); however, the opposite effect has also been reported (Szentirmai and Krueger, 2006; Ushimura et al., 2015). *Drosophila* NPF exerts similar effect on sleep: overexpression of NPF promotes sleep but is restricted to males (He et al., 2013); conversely, activation of NPF-producing cells promotes wakefulness (Chung et al., 2017). One possible reason for such contradictory effects is that the directions of sleep regulation by NPF and NPY may differ with the target neurons (Singh et al., 2017).

NPF is expressed in the brain and the midgut. NPF produced from the midgut is known to regulate germline stem cell proliferation, although its effect on sleep is unknown (Ameku et al., 2018). Furthermore, the relationship between NPF in the brain and sleep is not understood. *Drosophila* sleep is controlled by two oscillators: a morning oscillator and an evening oscillator. Dorsal lateral neurons (LNds) and fifth small ventral lateral neurons (sLNv) expressing NPF are known to be part of the evening oscillator, and their neural activity increases in the evening (Liang et al., 2016). LNds are composed of six neurons, three of which are NPF-positive neurons and one among which is a cryptocrome (CRY)-positive neuron (Yoshii et al., 2008). According to Chung *et al*., CRY-positive NPF-producing cells promotes wakefulness (Chung et al., 2017). However, the target neurons expressing NPFR are still not clear. Identification of NPF-producing cells that are controlled by the active substances of SBT2227 and the NPFR-expressing cells that receive its signal should be identified in future to understand the mechanism of action of SBT2227. This may also clarify the relationship between NPF/NPFR signals and sleep.

In our RNA-seq analysis, peptidoglycan muralytic activity, a GO category associated with bacterial recognition, was detected as an enriched category in the gut of flies treated with SBT2227. The immune-active state of animals enhances sleep, which has been reported in both mammals (Krueger et al., 1982) and flies (Kuo et al., 2010). SBT2227 harbors peptidoglycan on the cell wall surface, which is recognized by the Toll signaling pathway for gut immune responses (Valanne et al., 2011). Therefore, the cell wall fraction should be a potential substance for sleep promotion via immune responses. However, the precipitated fraction of crushed-SBT2227, which could be rich in peptidoglycan components, did not significantly affect fly sleep (**Figure 5**). Hence, the immune system may not be directly regulating SBT2227’s effects on sleep.

In conclusion, we found a novel biogenic action of *L. plantarum* strain SBT2227 as sleep-promoting effects on *D. melanogaster*. The active substance derived from cellular components of SBT2227 would be ingested, digested and absorbed via intestinal tract. The active substance could then act on the responsible brain circuits via the neuropeptide NPF to exert the sleep-promoting effect **(Figure 8**). Mammals also harbor the NPF homolog, NPY, which is reported to control sleep behavior. Therefore, we expected SBT2227 to exert sleep-promoting effects in mammals. Currently, the mainstream approach for improving sleep involves the use of pharmaceuticals. In the near future, elucidation of the mechanism of action of functional LABs that act on brain functions, such as SBT2227, will improve our understanding regarding new aspects of brain–gut– bacteria interactions and also assist in the development of functional fermented foods.

**Figure 8.**
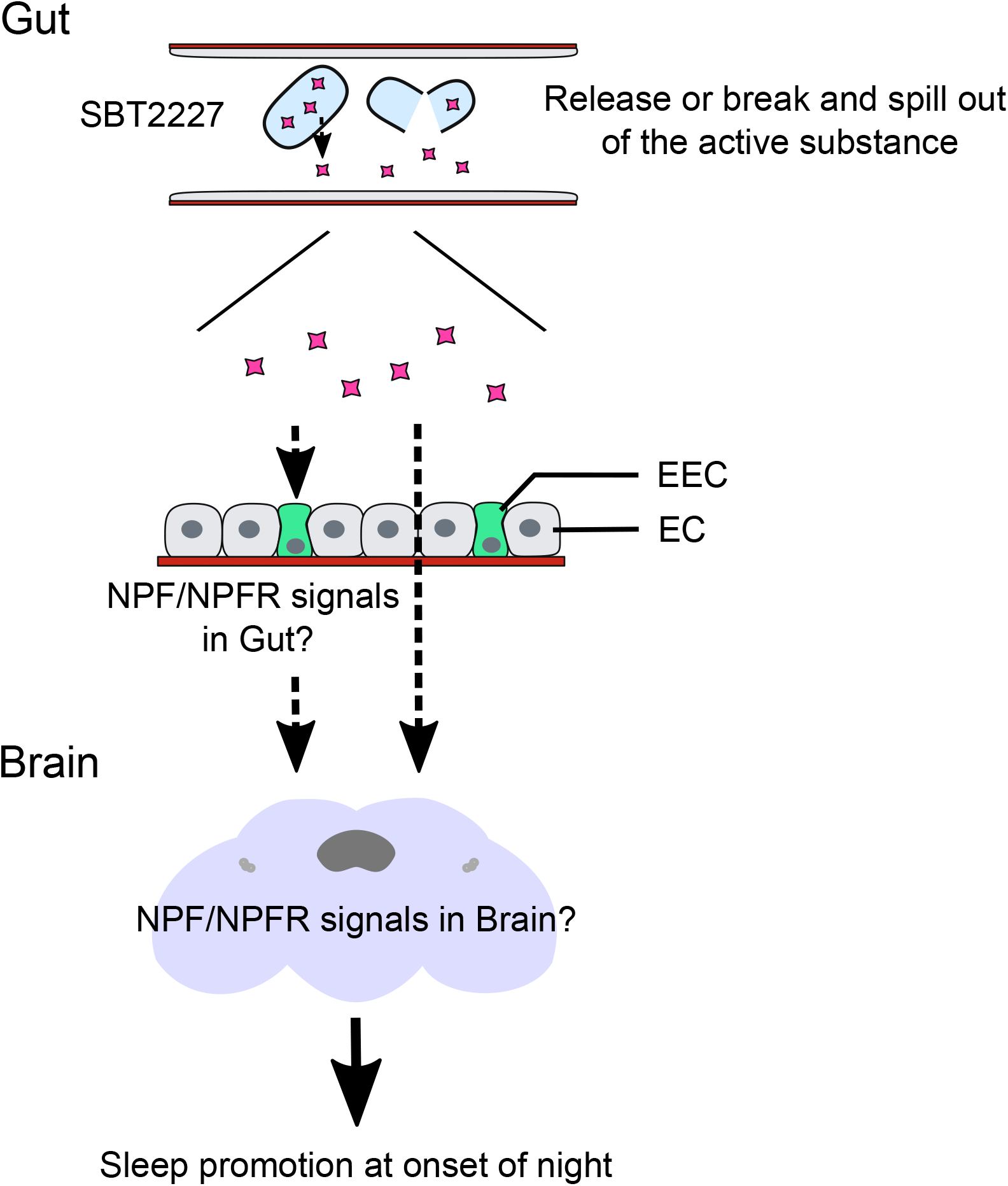
Model showing the biogenic action of the SBT2227 on fly sleep. The active substance of SBT2227 is released from the cells, or the cells are digested and flow out into the gut tract. The active substance acts directly or indirectly on NPF-producing cells and finally acts on the brain neuronal circuits to exert the sleep-promoting effect.

## LIMITATIONS OF THE STUDY

A contract with Megmilk Snow Bland Co., Ltd. is required for the use of *L. plantarum* SBT2227.

## AUTHOR CONTRIBUTIONS

T.K., H.M., and H.I. designed research; T.K., H.M., and H.I. performed research; H.I. contributed new reagents/analytic tools; T.K., H.M., and H.I. analyzed data; T.K. and H.I. wrote the paper; and A.K. and H.I. reviewed and edit the paper. All authors read and approved the manuscript.

## ACKNOWLEDGMENTS

We thank Dr. T. Awasaki, Dr. S. D. Pletcher, Dr. J. Y. Kwon, Bloomington Drosophila Stock Center, and Vienna Drosophila Resource Center, Kyoto stock center for fly stocks. We also thank Dr. I. Mori and Dr. T. Tanimura for comments and discussion, Y. Arai and Y. Ishikawa for fly maintenance. This work was supported by Grant-in Aid for Scientific research (C) (15K07147 and 18K06332 to HI), Inamori Foundation Research Grant, Japan to HI., and Research donation from Megmilk Snow Bland Co., Ltd.

## Lead Contact

Further information and requests for resources should be directed to and will be fulfilled by the lead contact Hiroshi Ishimoto.

## Materials Availability

*L. plantarum* SBT2227 was deposited in the International Patent Organism Depositary, National Institute of Technology and Evaluation (Chiba, Japan) under the accession number FERM BP-03104.

## Data and Code Availability

The original/source data are available from the lead contact on request.

## EXPERIMENTAL MODEL DETAILS

### Fly strains and rearing conditions

All fly strains used in this study were reared and maintained on standard cornmeal yeast (50 g/L glucose, 45 g/L yeast, 40 g/L corn flour, 8 g/L agar, 4 ml/L propionic acid, and 3 ml/L methyl 4-hydroxybenzoate) food at 24 ±1 °C and 60 ± 3 % humidity, with a 12 h light/12 h dark (12 h L/D) cycle. The *Canton-S*^*2202u*^ strain was used as the wild-type strain. *Amherst_3* (BDSC_4265), another wild-type strain, was used to confirm the sleep phenotypes obtained using *Canton-S*^*2202u*^. To suppress gene expression, the following *Gal4* and *UAS* strains were used: *Act-Gal4*^*25FO1*^ */ CyO, Act-GFP* (Dr. Awasaki, Kyorin Univ. Japan, RRID: BDSC_4414), and *tub5-GS-Gal4* (Scott D. Pletcher, Baylor College of Medicine, Houston, TX, USA). *UAS-AstA-RNAi* (BDSC_25866), *UAS-Mip-RNAi* (BDSC_41680), *UAS-AstC-RNAi* (BDSC_25868), *UAS-CCHa1-RNAi* (BDSC_57562), *UAS-CCHa2-RNAi* (BDSC_57183), and *UAS-Dh31-RNAi* (BDSC_41957) were obtained from BDRC. *UAS-NPF-RNAi* (VDRC ID: 108772) was obtained from the Vienna *Drosophila* Resource Center. For all experiments, virgin female flies were collected under CO_2_ anesthesia and used for experiments. Flies were maintained in food vials (< 20 flies per vial) until the experiments.

### Preparation of LAB samples

*L. plantarum* SBT2227 was obtained from Megmilk Snow Brand Co., Ltd. (Tokyo, Japan). SBT2227 was grown in Man, Rogosa, and Sharpe (MRS) broth (BD Biosciences, CA, USA) at 37°C for 16 h. The cultured bacteria were collected via centrifugation (7,000 × *g*, 20 min, 4°C) and then washed twice with sterilized saline (0.9% NaCl). The bacterial pellet was then suspended in a trehalose solution (12.5%) and concentrated to final volume of one-tenth of the culture medium. The bacterial suspension was incubated at 4°C for 2−5 h and then rapidly frozen in liquid nitrogen for storage at −80°C until further use. The frozen stock contained at least 5.2 × 10^10^ cfu/mL viable cells. For heat denaturation, SBT2227 was incubated at 65°C for 1 h using an incubator HB-80 (TAITEC Co., Saitama, Japan) or autoclaved at 121°C for 15 min using Panasonic MLS-3751 (PHC Corporation, Tokyo, Japan). After heat denaturation, the number of viable cells was <3,000 cfu/mL (65°C for 1 h) or <100 cfu/mL (autoclave). To crush SBT2227, 60.3 mg cells were suspended in 3 mL ultrapure water and placed in a 5.0 mL tube (TOMY SEIKO Co., Tokyo, Japan) along with 2 g of 0.1 mm glass beads (TOMY SEIKO Co., Tokyo, Japan). Cells were crushed at 4,200 rpm for 12 cycles at intervals of 30 s using a bead cell disrupter (TOMY SEIKO Co., Tokyo, Japan). After crushing, the glass beads were removed using a cell strainer (mesh size 40 am, Thermo Fisher Scientific, MA, USA).

## METHODS DETAILS

### Antibiotic treatment

The antibiotic-supplemented food contained 50 μg/mL tetracycline, 50 μg/mL ampicillin, 50 μg/mL kanamycin, and 15 μg/mL erythromycin in standard fly food. Newly enclosed flies were grown on antibiotic-supplemented food for four days. On the fourth day, the internal bacterial load was counted.

### Quantification of bacterial load in flies

Flies were washed in 70% ethanol for 15 s and then rinsed twice with sterile phosphate-buffered saline (PBS) for 15 s. Three flies were placed in 100 μL sterile PBS in a 1.5 mL tube and then homogenized using a sterilized pestle. The homogenates were diluted and plated on MRS agar plates. After 3 days of incubation under either aerobic or anaerobic conditions at 37°C, the number of colonies on the plate was counted. Anaerobic conditions were generated using AnaeroPack-Anaero (Mitsubishi Gas Chemical, Tokyo, Japan).

### Gram staining of bacteria

The bacterial suspension was spread on a glass slide, fixed with methanol, primary stained with Victoria blue, decolorized with picric acid in ethanol, and counterstained with fuchsin (Muto Pure Chemicals Co., Ltd. Tokyo, Japan).

### Sleep analysis

Fly sleep was measured using the *Drosophila* activity monitoring system (Trikinetics, MA, USA). Four-day old virgin female flies were transferred into a glass tube (5 × 65 mm) with control food or test food on one end of the tube, which was sealed with a cotton plug on the other end after loading the fly. The flies were allowed to acclimate to the new environment overnight, and locomotor activity was measured for 3−4 days at 25 ± 1°C in a 12 h light/dark cycle.

The control food contained trehalose (6.25%), sucrose (5.0%), and bacto-agar (1.0%). The LAB food contained SBT2227 (50%) in the control food. For evaluating crushed SBT2227 (10.1 mg/mL final concentration) and its centrifuged supernatant (5.4 mg/mL final concentration) and precipitate (3.0 mg/mL final concentration), each of them was mixed with the control food lacking trehalose. For the *GeneSwitch* system, RU486 (500 μM final conc., Mifepristone, Sigma-Aldrich, MO, USA) was mixed with SBT2227 food.

Locomotor data were collected in 1-min bins, and inactivity for at least 5 min was defined as a sleep bout. Behavior parameters were analyzed using R package Rethomics (Geissmann et al., 2019). Dead animals were excluded from analyses.

### RNA isolation

Total RNA was extracted using RNAiso Plus (Takara Bio, Shiga, Japan) according to the manufacturer’s protocol. Five flies were placed in 500 μL RNAiso Plus reagent and thoroughly homogenized with a plastic pestle. Total RNA was dissolved in 100 µL RNase-free water.

For RNA sequencing, total RNA was extracted from the fly gut using the RNeasy mini kit (Qiagen, Hilden, Germany) according to the manufacturer’s protocol. The guts were collected between ZT12-14 4 days after the start of the sleep experiment. The guts (from the lower part of the crop to the upper part of the Malpighian tubules) were dissected and immediately immersed in 2-mercaptoethanol with Buffer RLT of the RNeasy mini kit. This procedure was repeated thrice independently.

### Real-time polymerase chain reaction (PCR)

First-strand cDNA was synthesized from total RNA using ReverTra Ace qPCR RT master mix with gDNA Remover (TOYOBO, Osaka, Japan). Real-time PCR was performed using THUNDERBIRD SYBR qPCR Mix (TOYOBO, Osaka, Japan). All steps were performed according to the manufacturer’s protocol. PCR amplification was performed using the following primers: *NPF* primers were designed using the FlyPrimerBank (Hu et al., 2013). Ribosomal protein 49 (*rp49*) was used as the internal control.

*rp49*-f; 5′-AGTATCTGATGCCCAACATCG-3′

*rp49*-r; 5′-CAATCTCCTTGCGCTTCTTG-3′

*AstA*-f; 5′-TTGCACCGCGTATCCTGTCT-3′

*AstA*-r; 5′-ATGCTATGGGCACGGGATGG-3′

*CCHa1*-f; 5′-CCCAAATCGATGCCGACAATG-3′

*CCHa1*-r; 5′-GCAATTGGCCTCGGAATGTT-3′

*Mip*-f; 5′-CTCTAGCACCTAGTCTCCACG-3′

*Mip*-r; 5′-GTTGCCATTTGGTATGTATTGATGT-3′

*NPF*-f; 5′-TCCGCGAAAGAACGATGTCA-3′

*NPF*-r; 5′-CTCCTCATTAAAACCGCGAGC-3′

### RNA sequencing analysis

RNA sequencing libraries were constructed and sequenced by Macrogen Inc. using the Illumina NovaSeq6000 system. Read fragments were mapped to the *D. melanogaster* genome NCBI GCF_000001215.4_Release_6_plus_ISO1_MT, and assembled using StringTie (Pertea et al., 2015). edgeR (Robinson et al., 2010) was used for comparison of RNA expression levels and the exact test for negative binomial distribution was applied. Among the genes expressed in both the control group and the SBT2227 administered group, the expression level of the genes was more than 2-fold, and *p* < 0.05 were considered for DEGs. For genes expressed in only one of the groups, those with *p* < 0.05, were included in DEGs. We performed GO enrichment analysis of these DEGs using g:Profiler (https://biit.cs.ut.ee/gprofiler/).

## QUANTIFICATION AND STATISTICAL ANALYSIS

All statistical analyses were performed using the R Studio version 1.0.136. (www.r-project.org). Sleep data tended not to show a normal distribution, and hence, Wilcoxon-Mann-Whitney tests or Dwass-Steele-Critchlow-Fligner tests for nonparametric statistics were applied. When the Wilcoxon-Mann-Whitney tests were repeated, the *p* values were adjusted using the Bonferroni method. Statistical significance was set at *p* < 0.05. The boxes in box and whisker plots represent the median and interquartile range (the distance between the first and third quartiles), and whiskers represent the highest and lowest data points, excluding any outliers. To evaluate the sleep-promoting effect, the effect size *r* was calculated from the *z* value of the Wilcoxon-Mann-Whitney test using the following formula: *r* = z /√ n.

## SUPPLEMENTAL INFORMATION

Supplemental Information can be found online at https://xxxxxxxxx

**Figure S1.**
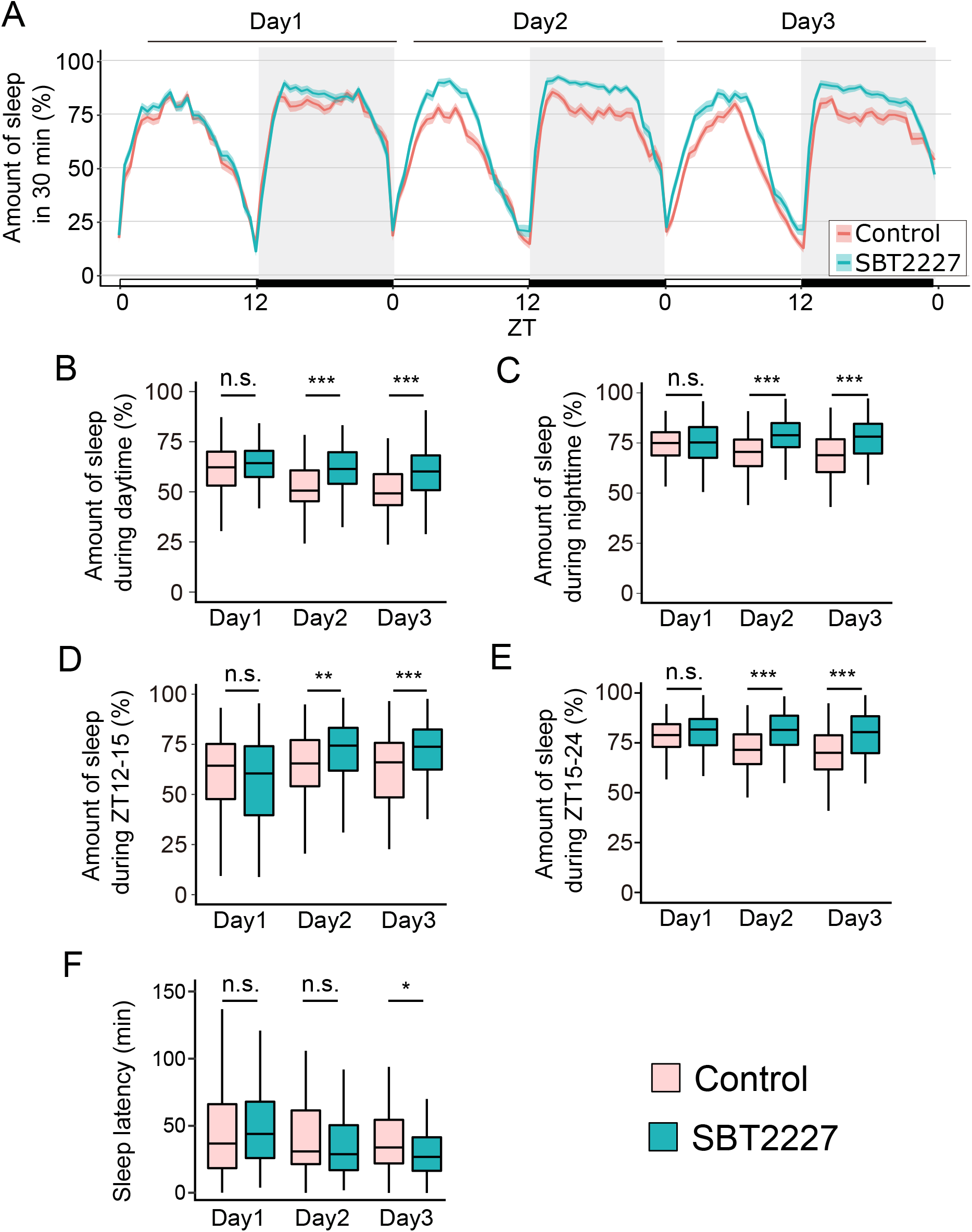
Sleep-promoting effects of SBT2227 on another wild-type strain, related to Figure 1. The sleep-promoting effects of SBT2227 were confirmed in another wild-type strain, *Amherst-3*, the genetic background of which differed from that of *Canton-S*^2202u^. (A) Sleep patterns of female *Amherst-3* fed the control food (red) or SBT2227 food (green). Sleep traces are presented as mean ± SEM. (B, C) Amount of sleep during daytime (ZT0-12) (B), amount of sleep during night-time (ZT12-15) (C). (D, E) Amount of sleep at specific timing. ZT12-15 (D) and ZT15-24 (E) are shown. (F) sleep latency. N = 96 for each group. The Wilcoxon-Mann-Whitney test was used for statistical analysis. * *p* < 0.05, ** *p* < 0.01; *** *p* < 0.001, n.s.; not significant.

**Figure S2.**
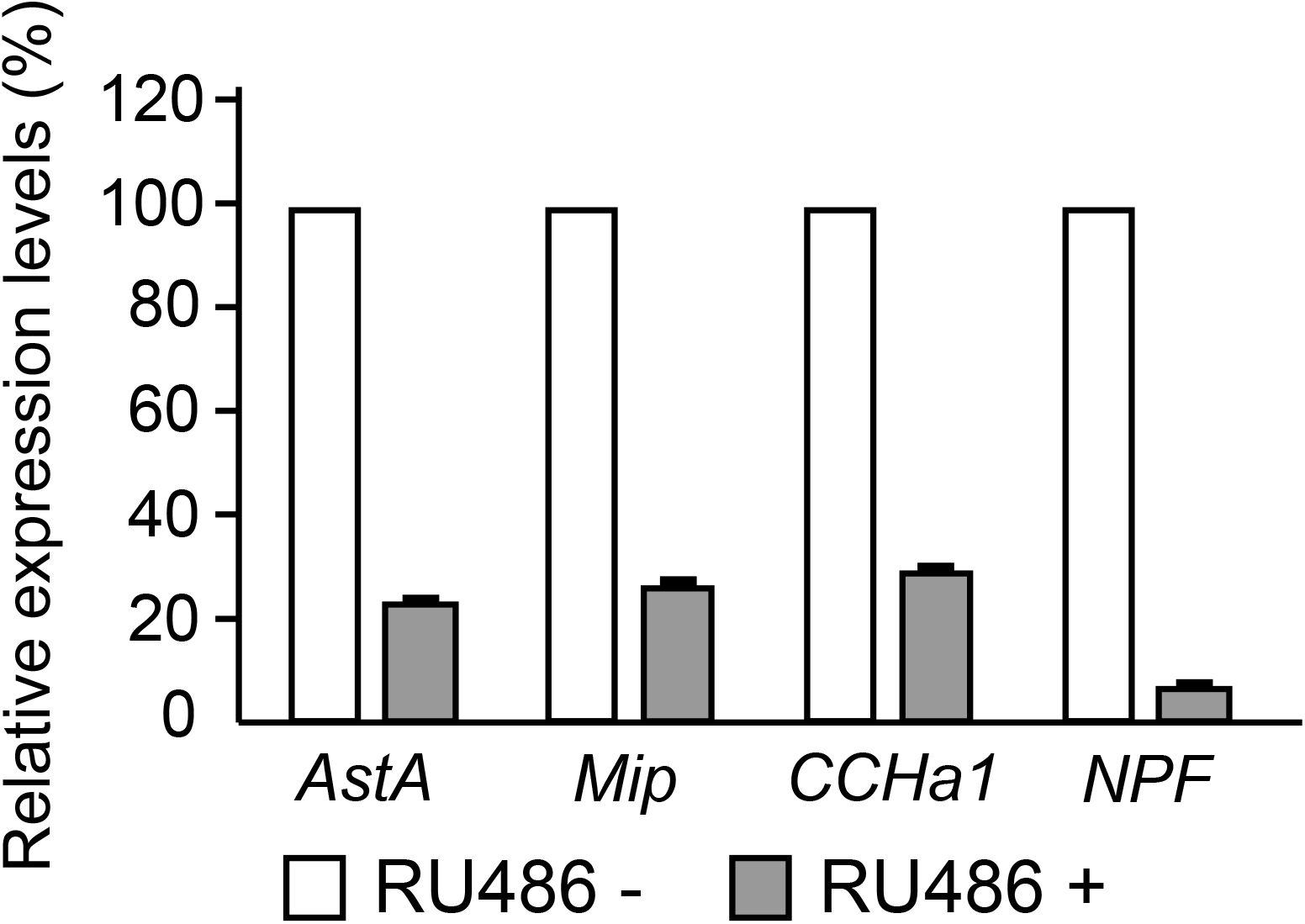
The peptide hormone genes were able to be knocked down by RNAi, related to figure 7. Efficiency of RNAi combined with *tub5-GS-Gal4*. Gene expression levels were measured on the second day after administration of RU486. n = 3, Bar graphs show mean and standard deviation.

**Table S1. A total of 787 genes were identified as differentially expressed genes (DEGs) by RNA-seq in the SBT2227-treated and non-treated groups, related to figure 6**.

